# Advancing Noninvasive Mechanical Ventilation: Simulating Techniques for Improved Respiratory Care

**DOI:** 10.1101/2024.10.01.616043

**Authors:** Md Toki Tahmid, Mrinmoy Nandi Bappa

## Abstract

Respiratory failure is a critical condition that often requires mechanical ventilation to support or restore normal breathing. While invasive mechanical ventilation (IMV) is commonly used for severe cases, noninvasive mechanical ventilation (NIMV) offers a less intrusive alternative that reduces complications and can be applied in moderate cases. The COVID-19 pandemic highlighted the global shortage of ventilators, particularly in low- and middle-income countries (LMICs), where limited access to life-saving equipment exacerbated the crisis. In response to these challenges, this paper presents a simplified, compartmental-based simulation model for NIMV. This model provides a practical and accessible tool for simulating respiratory system behavior under various ventilation modes, using the analogy between electrical circuits and lung physiology. By simulating key parameters such as airway resistance and lung compliance, the model allows clinicians and researchers to evaluate ventilator performance and optimize treatment strategies. Furthermore, the simulation offers a blueprint for developing cost-effective, easy-to-use NIMV systems that can be deployed in resource-constrained environments. Our contribution seeks to address the ventilator shortage by enabling better design and understanding of noninvasive ventilation, ultimately improving respiratory care for patients with moderate respiratory failure.

## 1 Introduction

Respiratory failure is a life-threatening condition that often necessitates the use of a mechanical ventilator (MV), a vital device that assists in breathing by facilitating the movement of air in and out of the lungs. Mechanical ventilation is typically classified as either invasive or noninvasive, both of which play crucial roles in critical care. Invasive mechanical ventilation, which involves the insertion of an endotracheal tube, is commonly used for severe respiratory conditions as it provides effective gas exchange and sputum drainage. However, invasive methods are often associated with complications such as ventilator-associated pneumonia, airway damage, and delirium, especially in low- and middle-income countries (LMICs) where healthcare resources are limited [11].

Noninvasive mechanical ventilation (NIMV), on the other hand, has emerged as a less intrusive alternative, particularly suited for patients with moderate respiratory failure. It delivers ventilatory support via a facial or nasal mask, eliminating the need for invasive procedures while still improving oxygen exchange and reducing the burden on fatigued respiratory muscles. NIMV can be controlled by either pressure or volume, depending on the patient’s needs. This method has proven effective in treating exacerbations of chronic obstructive pulmonary disease (COPD) and cardiogenic pulmonary edema, and its use has been widely recognized during the COVID-19 pandemic for managing acute hypoxemic respiratory failure [6, 4].

The COVID-19 pandemic, caused by the novel coronavirus SARS-CoV-2, was declared a global health emergency by the World Health Organization (WHO) in March 2020 [12]. It placed extraordinary pressure on healthcare systems across the globe, leading to a significant proportion of hospitalized patients developing severe respiratory issues that required intensive care unit (ICU) support and mechanical ventilation. The sudden surge in demand for ventilators resulted in severe shortages, particularly in LMICs, compounding the challenges of treating respiratory failure. The high cost of mechanical ventilators remains a major obstacle to providing adequate care, both for acute crises like COVID-19 and for chronic respiratory conditions. These shortages highlight the urgent need for cost-effective, accessible solutions [3].

In response to these challenges, modeling and simulating the respiratory system has become increasingly important in biomedical engineering. These simulations offer valuable insights into the complex dynamics of respiratory failure and mechanical ventilation, allowing researchers and clinicians to evaluate ventilator performance and test new ventilation strategies. Recent advancements in modeling have taken advantage of the analogy between electrical circuits and lung physiology, where resistors represent airway resistance and capacitors represent lung compliance. This approach provides a practical framework for simulating mechanical ventilation in both single- and multi-compartment lung models [5, 8]. For example, the Pressure-Controlled Ventilator (PCV) approach, applied to compartmental lung models, has been used to explore how different modes of ventilation affect respiratory behavior [7]. Tools like MATLAB’s Simulink enable researchers to simulate various mechanical ventilation modes, such as Volume-Controlled Ventilation (VCV), and to compare flow rate and pressure data against theoretical models for validation purposes [1, 10].

### Our Contribution

In this work, we contribute to the field by developing a straightforward, compartmental-based simulation model that can be used to easily simulate the behavior of the respiratory system under noninvasive mechanical ventilation. This model provides an accessible and practical tool for understanding how ventilators interact with the lungs, particularly for noninvasive ventilation scenarios. Using this simulation, clinicians and researchers can explore different ventilation modes and optimize them for patient care. Additionally, we demonstrate how this model can be implemented in real-world scenarios to guide the development of user-friendly, cost-effective noninvasive mechanical ventilators, which are essential in resource-limited settings.

In summary, mechanical ventilation is a critical intervention for patients with respiratory failure, and its importance has become even more pronounced during the COVID-19 pandemic. While invasive mechanical ventilation is the gold standard for severe cases, noninvasive methods offer a viable, less risky alternative. Our work focuses on addressing the global shortage of ventilators by offering an easy-to-use simulation model for noninvasive ventilation, which has the potential to inform the development of affordable and accessible ventilator solutions in resource-constrained environments.

## 2 Methodology

### 2.1 Overall Architecture

The proposed system simulates the effects of ventilation on the respiratory system using a Pressure Controlled Ventilation (PCV) model. This simulation framework is designed to assess how different ventilatory strategies and parameters influence respiratory mechanics, especially in various lung models (single, multi-series, and parallel-compartmental). The entire simulation was implemented in MATLAB/Simulink to ensure an efficient and reliable test platform before hardware implementation. By using this environment, different clinical scenarios can be simulated, and the system’s performance can be optimized.

#### System Design Overview

- The PCV model controls ventilation by generating pressure waveforms that replicate human breathing cycles.
- These pressure waveforms are applied to different mathematical lung models to simulate real-world ventilation conditions.
- Lung models are represented by electrical analogies, where pressure, airflow, and lung volume correspond to voltage, current, and charge respectively.
- Different modes of operation (time-triggered and pressure-triggered) are implemented to replicate spontaneous breathing as well as mechanical ventilation for patients who are unable to breathe unaided.

The architecture of the system includes two primary ventilator modes: *Time-Triggered Control Mode* and *Pressure-Triggered Spontaneous Breathing Mode*. Both are described in the following sections.

### 2.2 Simulating Ventilation Effects on the Respiratory System

Before moving to hardware implementation, the entire system was first modeled in a simulation environment to assess its performance, effectiveness, and ensure optimization. This included the development of a mathematical model (MM) of the Pressure Controlled Ventilator (PCV) for simulating the impact of pressure waveforms on single, multi-series, and parallel-compartment lung models [8]. All simulations were performed using MATLAB/Simulink. The ‘Continuous’ Powergui block was used for simulating electrical systems with continuous-time elements.

#### 2.2.1 Modeling the Ventilator Function

The ventilator operates in two modes: *Time-Triggered Control Mode* for cases when the patient is unable to breathe properly and requires mechanical ventilation, and *Pressure-Triggered Spontaneous Breathing Mode* for assisting patients who can initiate breaths.

##### Mode 1: Basic PCV -Time-Triggered Control

The first ventilator mode models a time-triggered breathing cycle, where a constant pressure waveform is generated to represent normal human breathing, independent of patient effort. The PCV generates a pressure waveform that consists of two phases: an inspiratory phase and an expiratory phase [8]. The pressure waveform is mathematically modeled as:

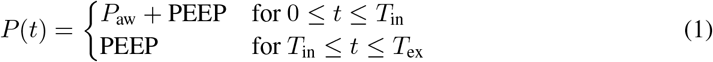

Here, *P* (*t*) represents the pressure signal of the PCV, *PEEP* is the positive end-expiratory pressure, and *P*_aw_ is the pressure in the respiratory airways. The total pressure waveform is a combination of the inspiratory pressure (IP) during the inspiratory phase (*T*_in_) and the expiratory pressure (EP), where the EP is assumed to be equal to PEEP during the expiratory phase (*T*_ex_).

The total cycle time (TCT) and respiratory rate (RR) are computed as:

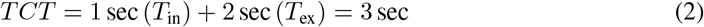

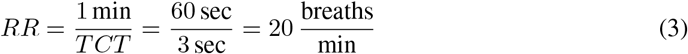

The respiratory rate (RR) and other ventilator parameters were selected based on normal adult ranges, as shown in Table 1.

**Table 1:**
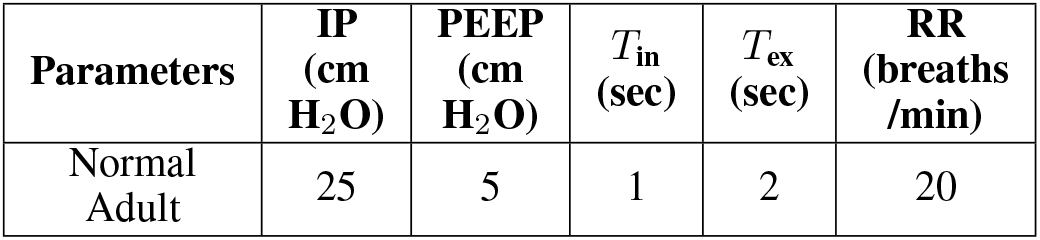
Ventilation Parameter for Normal Adult.

##### Mode 2: Pressure-Triggered Spontaneous Breathing

In this mode, a pressure sensor placed inside the patient’s mask monitors the pressure *P*_*s*_ in the airways. If *P*_*s*_ exceeds an upper threshold, the ventilator detects that the patient is exhaling, and the pressure signal *P* (*t*) is set to PEEP. If *P*_*s*_ falls below a lower threshold, the ventilator detects that the patient is inhaling, and *P* (*t*) is set to *P*_aw_ + PEEP. Otherwise, *P* (*t*) remains unchanged. This mode adapts to the patient’s breathing cycle, meaning that the inspiratory and expiratory times are not fixed.

#### 2.2.2 Modeling the Respiratory System

In this work, the respiratory system is modeled as an electrical circuit, where lung compliance and resistance are represented by capacitance and resistance respectively. Three versions of the lung model are simulated: a one-compartmental model, a series two-compartmental model, and a parallel two-compartmental model [5, 8].

##### One Compartmental Model

The *One Compartmental Model* simplifies the lung’s mechanical behavior by modeling it as a single electrical circuit. In this analogy, the lung’s resistance to airflow is represented by a resistor *R*, and its ability to expand and contract (compliance) is represented by a capacitor *C*. The electrical analogy works because airflow and pressure in the lungs follow similar principles to current and voltage in an electrical circuit.

The *transfer function* of the system is given by the equation:

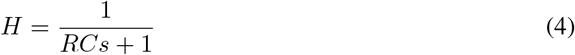

This equation describes the relationship between pressure and flow in the lung. Specifically: -*R* represents the *resistance* of the airways to airflow, where *R* = 2 cm H_2_O/L/s. -*C* represents the *compliance* of the lung, which is a measure of its ability to expand in response to pressure, where *C* = 0.1 L/cm H_2_O.

The product of *R* and *C* determines the *impedance Z* of the lung system. This impedance affects how air flows into and out of the lungs, dictating the lung’s inflation and deflation during each breathing cycle. Lower impedance allows for easier breathing, while higher impedance indicates stiffer or more resistant lungs, requiring greater effort to achieve the same airflow.

This model serves as a fundamental starting point for understanding lung mechanics in health and disease, providing insights into conditions like asthma and chronic obstructive pulmonary disease (COPD), where airway resistance is altered.

##### Series Two Compartmental Model

The *Series Two Compartmental Model* provides a more detailed representation of lung function by modeling the lung as two distinct compartments connected in series. This model introduces the concept of spatial variation within the lung, allowing for a more accurate description of airflow and pressure distribution between different parts of the respiratory system.

The transfer function for this system is given by:

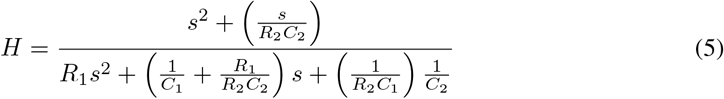

In this equation: -*R*_1_ and *R*_2_ represent the *resistances* of two different lung compartments. -*C*_1_ and *C*_2_ represent the *compliances* of those compartments. -The series connection implies that airflow must pass through the first compartment to reach the second, and any changes in the parameters of one compartment directly affect the overall system’s behavior.

This model is particularly useful for understanding lung conditions where different regions of the lung exhibit varying degrees of resistance or compliance. For instance, in diseases such as emphysema or fibrosis, different regions of the lung may behave differently, leading to uneven airflow distribution and varying levels of gas exchange efficiency.

##### Parallel Two Compartmental Model

In this model, the lung is represented by two parallel compartments (Figure 3). The **Parallel Two Compartmental Model** allows for a more realistic simulation of lung mechanics when both compartments function independently but contribute simultaneously to overall lung volume and airflow.

**Figure 1.**
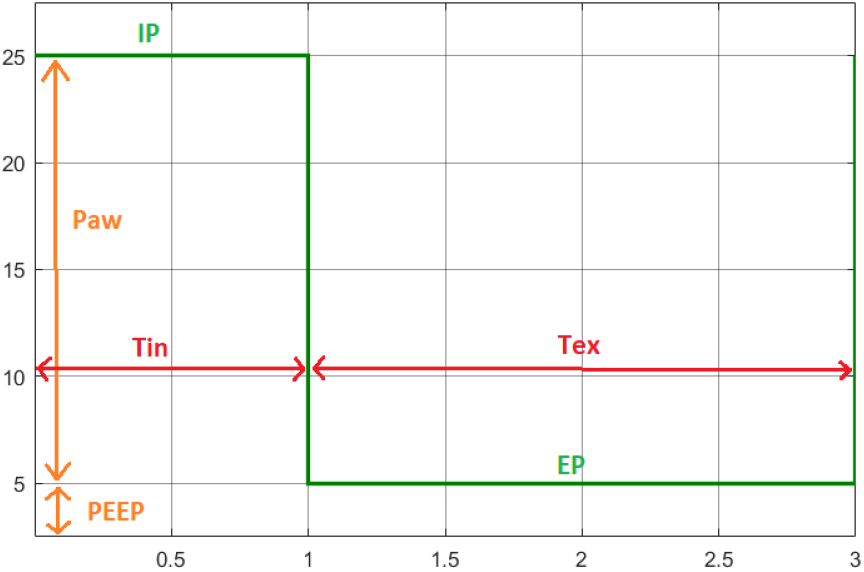
Pressure waveform of PCV signal during normal ventilation

**Figure 2.**
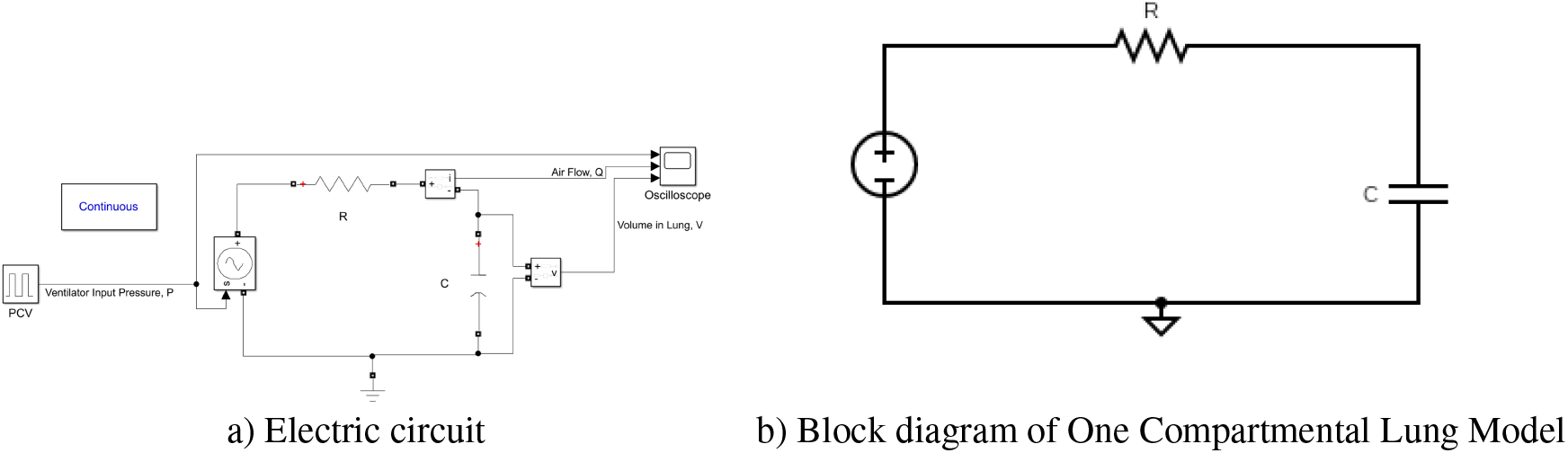
a) Electric circuit, (b) Block diagram of One Compartmental Lung Model

**Figure 3.**
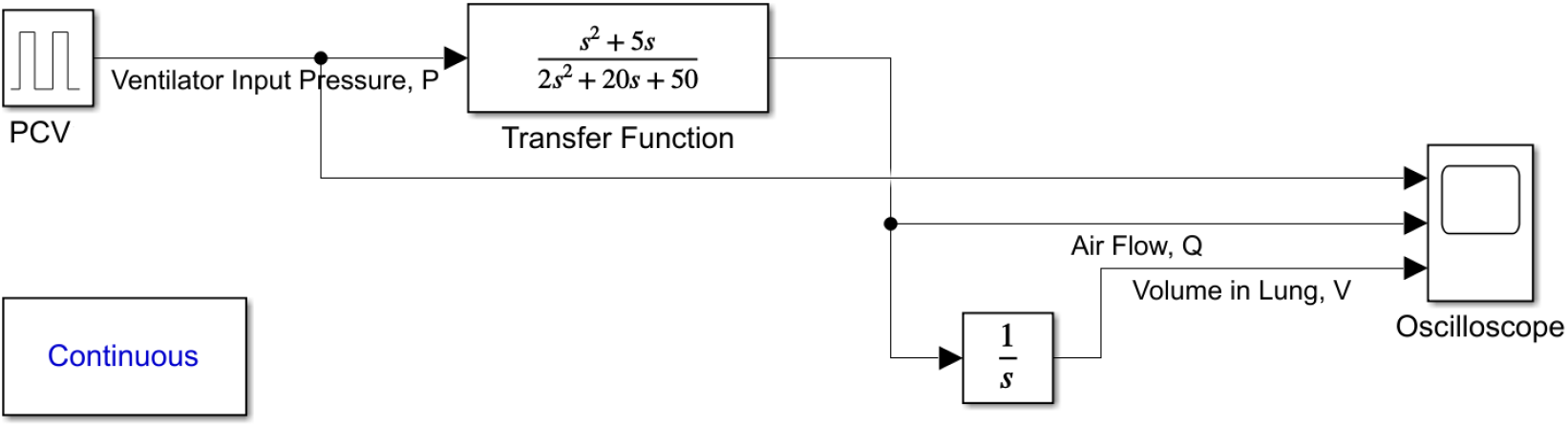
Electric circuit representation for two compartmental model

The transfer function for this model is given by:

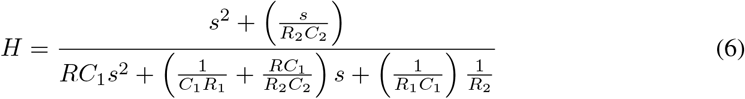

Here, *R*_1_ and *R*_2_ again represent the resistances of the two compartments, while *C*_1_ and *C*_2_ represent their compliances. The parallel connection allows each compartment to function somewhat independently, simulating real-life conditions where parts of the lung can have different resistances or compliances due to localized pathologies such as in obstructive lung diseases.

This model is particularly valuable in understanding conditions like acute respiratory distress syndrome (ARDS) or pneumonia, where different parts of the lung may be affected differently by the disease process. The parallel model helps predict how various treatments, such as mechanical ventilation, will impact different parts of the lung.

## 3 Results and Discussion

This section presents the results of simulating the Pressure Controlled Ventilation (PCV) model applied to different lung models under both normal and abnormal conditions. The analysis includes comparisons of lung responses (pressure, volume, and flow rate) across multiple compartmental models, examining both normal and diseased respiratory conditions.

### 3.1 Simulation Results and Analysis

#### 3.1.1 Comparison of Lung Model Results

The results of applying the PCV signal to various mathematical models (MMs) of the lungs, including the one-compartmental, series two-compartmental, and parallel two-compartmental models, are displayed in Fig.4. These models simulate lung dynamics in terms of instantaneous pressure (*P*), lung volume (*V*), and airflow rate (*Q*).

**Figure 4.**
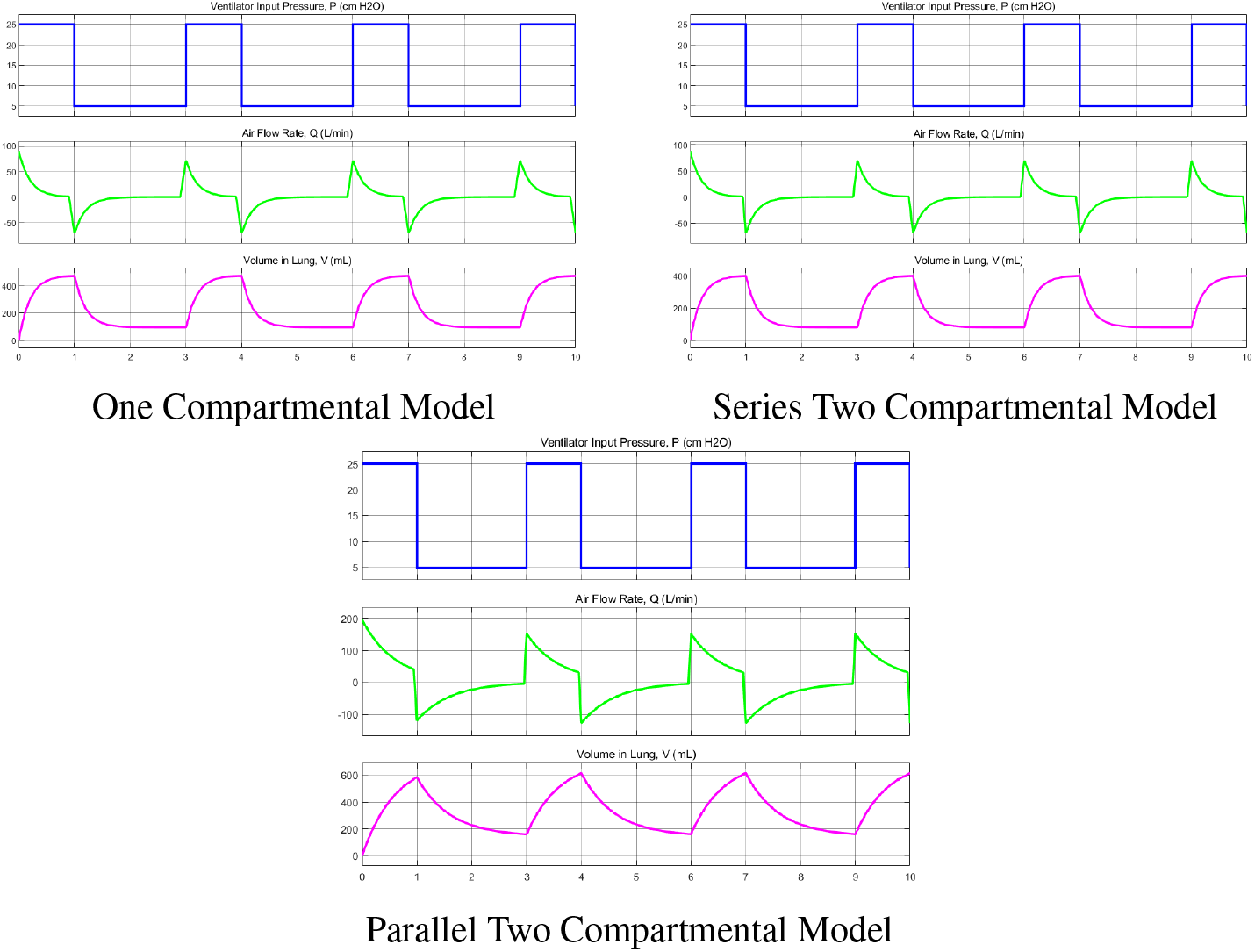
Comparison of Pressure, Flow, and Volume for Different Compartmental Models

As shown in Fig.4, the pressure waveform and flow rate are cyclical, reflecting the alternating phases of inspiration and expiration. The flow rate returns to zero at the end of the inspiratory cycle and assumes an inverse direction during the expiratory phase. Both the inspiration and expiration cycles maintain consistent durations, resulting in symmetrical waveforms. Typically, the flow rate at the beginning of inspiration is higher compared to the start of expiration. The total respiration time was chosen as 9 seconds (3 full cycles).

Lung volume increases during inspiration and decreases during expiration. The maximum volume reached is approximately 500 mL, which is consistent with normal adult tidal volumes [2, 9]. The pressure and volume waveforms provide valuable insight into lung mechanics. For example, the increased resistance in the lungs (modeled in later sections) would alter the slope of the pressure and volume curves, slowing the filling and emptying of the lungs.

Among the three compartmental models, the results of the parallel two-compartmental model show significant deviations. The parallel model produces deformed shapes in the pressure, volume, and flow rate waveforms, along with larger amplitude variations that exceed normal physiological ranges. These deviations suggest a more unstable or inconsistent mechanical response in comparison to the single-compartmental or series models. Consequently, for further investigation and optimization, the one-compartmental model, which provides a more stable and realistic response, is selected as the baseline model.

In conclusion, the simulation results indicate that the one compartmental and series two compartmental models yield stable, physiologically realistic outputs, while the parallel two-compartmental model introduces distortions and abnormal amplitudes. For further simulations and optimizations, the one-compartmental model is chosen as the most reliable model.

#### 3.1.2 Comparison of PCV Signal Variations

In addition to the standard PCV signal, an anomalous case was simulated to investigate the effect of changes in ventilator settings. As shown in Table 2, the abnormal case increases the inspiratory pressure (IP) and positive end-expiratory pressure (PEEP), which impacts the lung volume and flow rate.

**Table 2:**
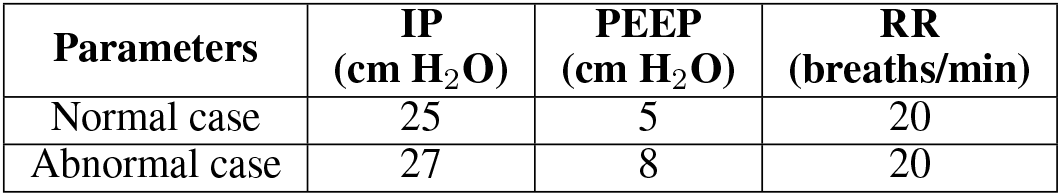
Parameter values of the PCV signal in normal and abnormal cases.

| Parameters | IP<br>(cm H<sub>2</sub>O) | PEEP<br>(cm H<sub>2</sub>O) | RR<br>(breaths/min) |
| --- | --- | --- | --- |
| Normal case | 25 | 5 | 20 |
| Abnormal case | 27 | 8 | 20 |

Fig. 5 displays the results of applying the abnormal PCV signal to the one-compartmental lung model. While the pressure and flow rate waveforms remain similar in shape to the normal case, there is a noticeable increase in lung volume. The lung volume reaches approximately 550 mL, compared to the normal 500 mL, suggesting that the elevated inspiratory pressure results in greater lung inflation. This deviation could potentially cause overinflation and barotrauma in patients, particularly those with compromised lung function.

These results suggest that careful adjustment of inspiratory pressure and PEEP settings is critical for preventing overinflation and maintaining lung safety during ventilation. The analysis demonstrates that while small increases in these parameters may have minor effects on flow rate and pressure, lung volume can increase significantly, leading to potential complications.

#### 3.1.3 Comparison of Lung Resistance in Diseased Conditions

To further evaluate the performance of the ventilator in diseased conditions, the effect of increased lung resistance was simulated. Obstructive pulmonary diseases, characterized by narrowing or blockage of the airways, were modeled by raising the resistance *R* of the lung. The higher resistance impairs lung expansion and reduces airflow, as demonstrated in Fig. 6.

As shown in Fig. 5, increased lung resistance reduces the lung’s ability to expand, leading to smaller volumes during the inspiratory phase. The rate of lung inflation also slows down, as indicated by the shallower slope of the volume curve. Additionally, airflow variations (Fig. 6) demonstrate a decrease in peak inspiratory flow and a delayed return to baseline during expiration.

**Figure 5.**
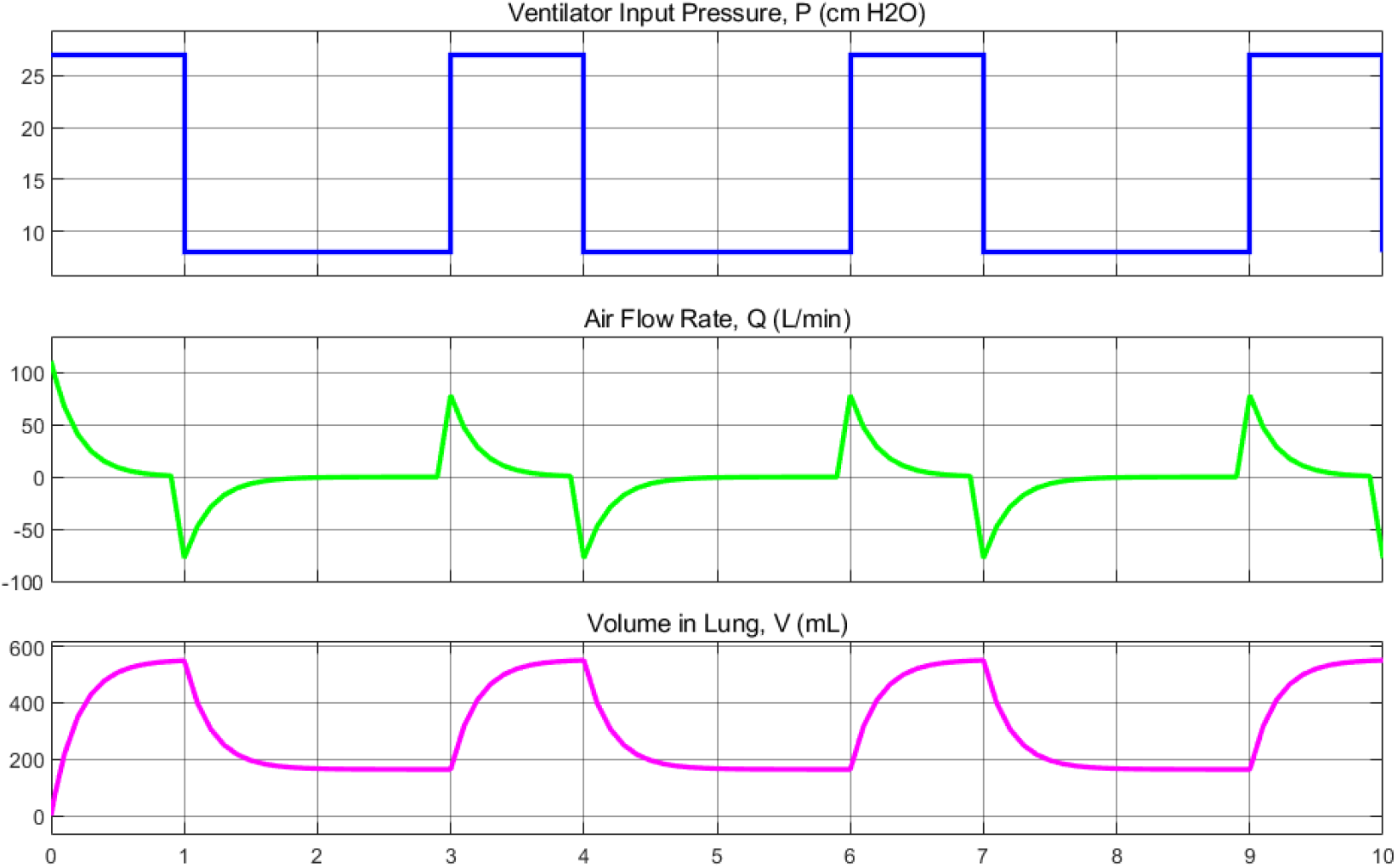
Air Flow, Pressure, and Volume Response in Abnormal Condition

**Figure 6.**
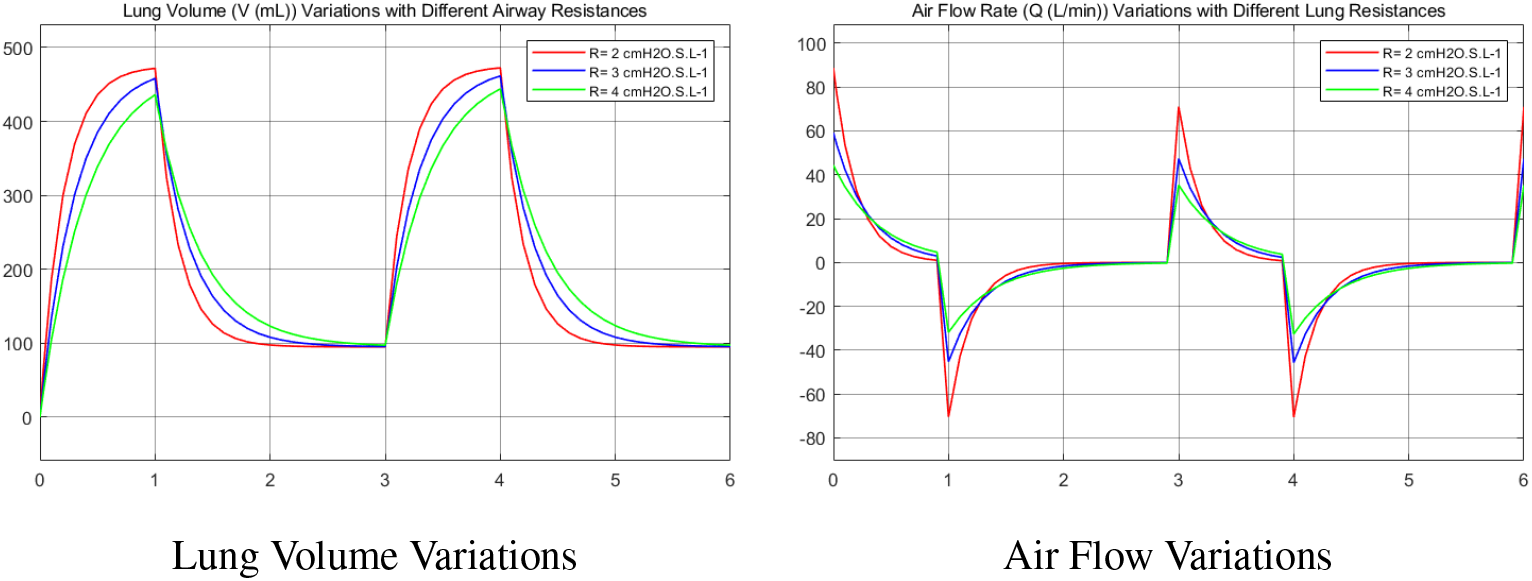
Comparison of Lung Volume and Air Flow Rate with Varying Lung Resistances

These results highlight the challenges of managing patients with obstructive pulmonary diseases. The ventilator must compensate for the increased resistance by adjusting the inspiratory pressure or extending the inspiratory time, to ensure adequate lung expansion and gas exchange.

In conclusion, the simulation of diseased conditions, particularly increased lung resistance, reveals the importance of dynamic ventilator settings in addressing abnormal lung mechanics. By tuning parameters such as inspiratory pressure, PEEP, and inspiratory time, the ventilator can ensure effective ventilation even in compromised patients.

## 4 Conclusion

In this work, we have developed a compartmental-based simulation model to aid in the understanding and optimization of noninvasive mechanical ventilation (NIMV) for patients with moderate respiratory failure. This model provides a straightforward and accessible tool for simulating key physiological parameters such as airway resistance and lung compliance, helping to evaluate the performance of NIMV systems under various scenarios. By leveraging this model, healthcare providers and researchers can better understand how noninvasive ventilators interact with the respiratory system, which is especially important given the high demand for ventilators and limited resources in many parts of the world.

The simulation can also inform the development of affordable, easy-to-use NIMV devices that are crucial in low- and middle-income countries, where ventilator shortages are most acute. Our work demonstrates how computational models can play a key role in addressing global health challenges by providing a platform for innovation and practical solutions. Moving forward, this approach can be expanded to further enhance the design and testing of noninvasive ventilators, contributing to better respiratory care worldwide.

## Notes

### Competing Interest Statement

The authors have declared no competing interest.

